# Interplay between intrinsic reprogramming potential and microenvironment controls neuroblastoma cell plasticity and identity

**DOI:** 10.1101/2021.01.07.425710

**Authors:** Cécile Thirant, Agathe Peltier, Simon Durand, Amira Kramdi, Caroline Louis-Brennetot, Cécile Pierre-Eugène, Ana Costa, Amandine Grelier, Sakina Zaïdi, Nadège Gruel, Irène Jimenez, Eve Lapouble, Gaëlle Pierron, Hervé J. Brisse, Arnaud Gauthier, Paul Fréneaux, Sandrine Grossetête-Lalami, Laura G. Baudrin, Virginie Raynal, Sylvain Baulande, Angela Bellini, Jaydutt Bhalshankar, Angel M. Carcaboso, Birgit Geoerger, Hermann Rohrer, Didier Surdez, Valentina Boeva, Gudrun Schleiermacher, Olivier Delattre, Isabelle Janoueix-Lerosey

## Abstract

Two cell identities, noradrenergic and mesenchymal, have been characterized in neuroblastoma cell lines according to their epigenetic landscapes relying on specific circuitries of transcription factors. Yet, their relationship and relative contribution in patient tumors remain poorly defined. Here, we demonstrate that the knock-out of *GATA3*, but not of *PHOX2A* or *PHOX2B*, in noradrenergic cells induces a mesenchymal phenotype. Our results document spontaneous plasticity in several models between both identities and show that plasticity relies on epigenetic reprogramming. We demonstrate that an *in vivo* microenvironment provides a powerful pressure towards a noradrenergic identity for these models. Consistently, tumor cells with a mesenchymal identity are not detected in a series of PDX models. Further study of the intra-tumor noradrenergic heterogeneity reveals two distinct cell populations exhibiting features of chromaffin-like or sympathoblast-like cells. This work emphasizes that both external cues of the environment and intrinsic factors control plasticity and cell identity in neuroblastoma.

## INTRODUCTION

Neuroblastoma is a childhood cancer arising from the peripheral sympathetic nervous system, known to be derived from multipotent neural crest cells (NCCs). Tumors mostly develop in the adrenal gland but a subset of them originates from sympathetic ganglia along the paravertebral sympathetic chain^1^. With respect to these localizations, neuroblastoma likely arises from the transformation of sympathoblasts either in sympathetic ganglia or in the adrenal medulla, or from catecholamine-secreting chromaffin cells of the adrenal medulla, or alternatively from a common sympatho-adrenal progenitor^2,3^.

The hallmark of neuroblastoma is its wide range of clinical presentations and outcomes, ranging from spontaneous regression to fatal outcome despite multimodal therapies^1^. High-risk neuroblastoma most often initially responds to intensive chemotherapy; however, relapses frequently occur followed by fatal outcome. Several genes including *MYCN*^4^, *ALK*^5–8^ and *TERT*^9–11^ have been identified as key drivers of neuroblastoma oncogenesis.

The master transcriptional regulators controlling the gene expression program of neuroblastoma have been recently highlighted through the characterization of the neuroblastoma super-enhancer landscape of neuroblastoma cell lines, revealing two distinct cell identities: a sympathetic noradrenergic identity defined by a core regulatory circuitry (CRC) module including the PHOX2B, HAND2 and GATA3 transcription factors (TFs) and a NCC-like/mesenchymal identity, close to that of human neural crest cells (NCCs), driven by TFs of the AP1 family among others^12,13^. Additional TFs participating in the noradrenergic CRC have been then identified, including ISL1, TBX2 and ASCL1^14–16^. It is likely that PHOX2A, also highly expressed in neuroblastoma cell lines and exhibiting the same DNA binding domain as PHOX2B is involved in the noradrenergic CRC. Importantly, mesenchymal tumor cells *in vitro* have been shown to be more resistant to standard chemotherapy^12,13^ suggesting that they may be involved in therapeutic resistance and relapses in neuroblastoma patients. Bulk RNA-seq analyses or immunohistochemistry with few markers suggested that mesenchymal tumor cells are present in patient tumors and that some tumors exhibit a mesenchymal identity^12,13,17^. Cellular plasticity between the noradrenergic and mesenchymal states has been reported for a few cell lines^13,18^, still the underlying mechanisms wherein the cell phenotype switches from one to the other state (considered as transdifferentiation) remain poorly described. In the present paper, we have combined various approaches including genetic inactivation of specific TFs and single-cell transcriptomic analyses to unravel TFs involved in cell identity and plasticity and better decipher the relationship and relative contribution of cells of noradrenergic and mesenchymal identity in neuroblastoma tumors.

## RESULTS

### The knock-out of *GATA3* but not of *PHOX2B* or *PHOX2A* induces a switch from a noradrenergic to a mesenchymal identity

In order to directly address the functional role of the PHOX2A, PHOX2B and GATA3 TFs in shaping the noradrenergic identity, we performed their individual genetic knock-out (KO) in the noradrenergic SH-SY5Y cell line using a CRISPR-Cas9 approach. Guide RNAs were designed to induce large deletions and create frameshift mutations in those genes, leading to truncated and nonfunctional proteins. We obtained two *PHOX2A^−/−^* clones, a *PHOX2B^−/−^* clone and two *GATA3^−/−^* clones (**Figure S1A**). While the *PHOX2A*^−/−^ and *PHOX2B*^−/−^ clones showed a morphology close to that of the parental noradrenergic SH-SY5Y cells with neurite-like processes, *GATA3*^−/−^ cells exhibited a more abundant cytoplasm and many actin stress fibers consistent with a mesenchymal phenotype (**Figure 1A**). *PHOX2A*^−/−^ and *PHOX2B^−/−^* clones maintained the expression of the noradrenergic CRC TFs including GATA3 and HAND2 (**Figures 1B and S1B**). Strikingly, both *GATA3^−/−^* clones showed an absence or highly reduced expression of the TFs from the noradrenergic CRC, including PHOX2A, PHOX2B, and HAND2 (**Figure 1B**). *PHOX2B^−/−^* and *GATA3^−/−^* cells were characterized by a decreased proliferation compared to the parental SH-SY5Y cell line (**Figure 1C**). Consistently with their phenotype, *GATA3^−/−^* clones displayed mesenchymal properties such as an increased invasion ability measured in a 3D-spheroid assay (**Figure 1D**), increased migration capacity documented using a transwell assay (**Figure 1E**) and a higher resistance to chemotherapy *in vitro* (**Figure 1F**). Bulk RNA-seq analysis showed that *PHOX2A*^−/−^ and *PHOX2B*^−/−^ clones harbored transcriptomic profiles highly similar to that of the parental noradrenergic SH-SY5Y cell line, confirming that *PHOX2A* or *PHOX2B* genetic invalidation did not change the cell identity. In contrast, *GATA3*^−/−^ clones, at two different passages called early and late showed a mesenchymal transcriptomic profile, as they clustered with the mesenchymal SH-EP cell line (**Figures 1G and S1C**). Accordingly, Gene Ontology analysis on the 2,938 differentially expressed genes between the transcriptomic profiles of the 4 *GATA3^−/−^* samples versus 5 noradrenergic neuroblastoma cell lines (without *MYCN* amplification as the parental SH-SY5Y) retrieved categories related to neuron differentiation, neurogenesis, sympathetic nervous development for the downregulated genes and extracellular matrix organization, cell motility and cell migration for the upregulated genes (**Figure S1D**).

**Figure 1.**
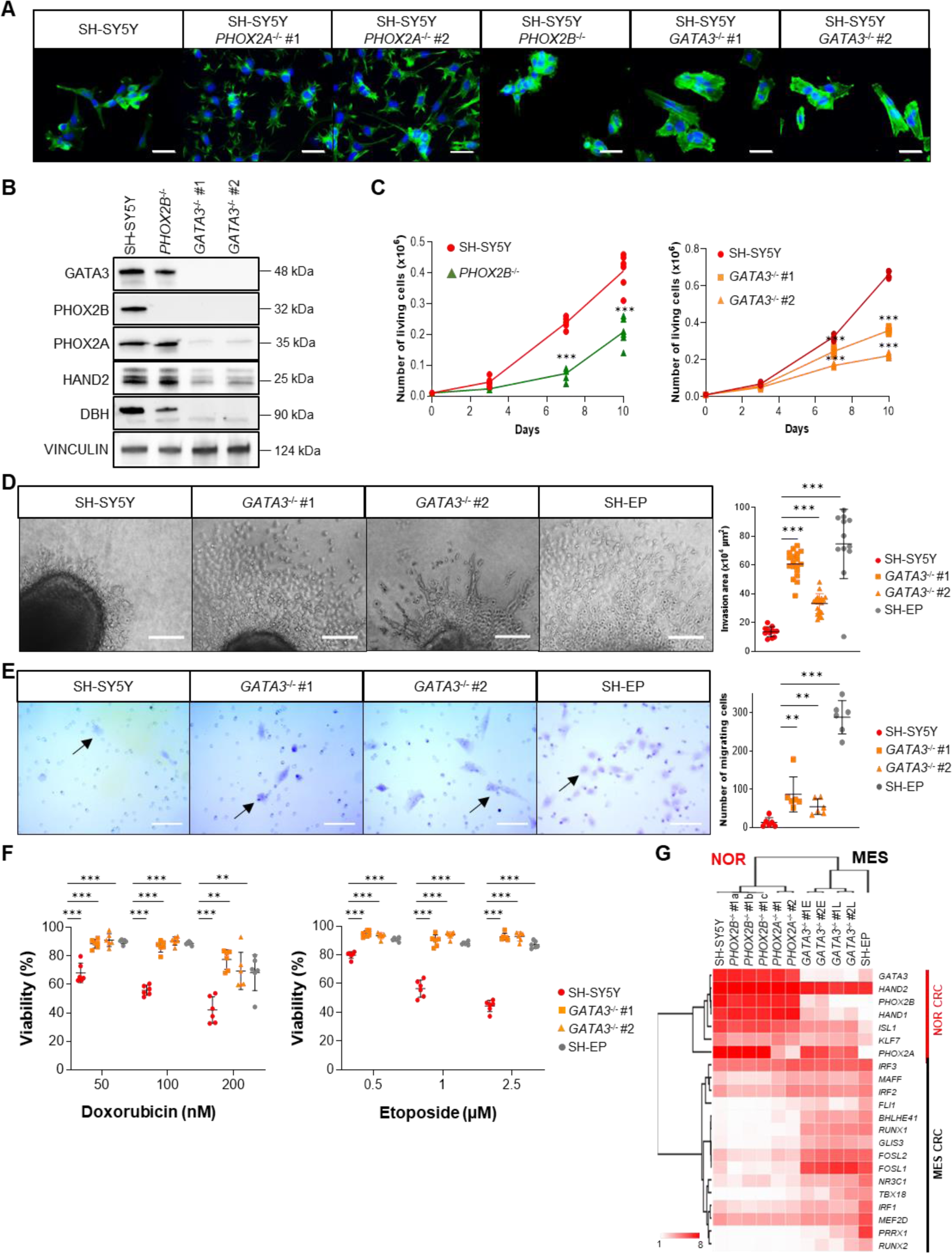
*GATA3^−/−^* cells exhibit mesenchymal properties whereas *PHOX2B^−/−^* or *PHOX2A^−/−^* cells keep a noradrenergic phenotype *in vitro.* (**A**) Phalloidin staining reveals the cell phenotype of the different KO clones and the SH-SY5Y parental cell line (x400, scale bar = 20 μm). (**B**) Western blot analysis of GATA3, PHOX2B, PHOX2A, HAND2 and DBH in the *PHOX2B*^−/−^ and *GATA3*^−/−^ clones, with Vinculin as a loading control. (**C**) Proliferation curves for *PHOX2B*^−/−^ and *GATA3* ^−/−^ cells *in vitro*. 10,000 cells were seeded in 24-well plates. Living cells were counted after 3, 7 and 10 days of culture (mean ± sd.; n = 6 replicates). (**D**) Representative images (left) and quantification (right) of spheroid invasion assays. Cells were seeded in low adherent plates to form neurospheres for 4 days and embedded in collagen I for 72h. Scale bar = 100 μm (mean ± sd.; n = 12 replicates for SH-SY5Y and SH-EP cell lines and n = 18 replicates for *GATA3*^−/−^ clones #1 and #2). (**E**) Representative images (left) and quantification (right) of transwell assays for the SH-SY5Y cells, the *GATA3^−/−^* clones (#1 and #2) and mesenchymal control cells (SH-EP). 50,000 cells were plated in a transwell insert and the living cells that have migrated into the transwell membrane were counted 24h later. Scale bar = 20 μm (mean ± sd.; n = 6 replicates). Each dot represents a replicate and is the mean of 6 different images of the well. (**F**) *GATA3*^−/−^ cells are more resistant to chemotherapy than the SH-SY5Y noradrenergic parental cell line. Cell viability was measured after 48 hours of chemotherapy treatments (Doxorubicin 50, 100, 200 nM and Etoposide 0.5, 1, 2.5 μM) (mean ± sd.; n = 6 replicates). (**G**) Heatmap showing the expression levels of the TFs of the noradrenergic (n=7) and mesenchymal (n=15) CRC in the *PHOX2B^−/−^* (clone#1, 3 replicates)*, PHOX2A^−/−^* (clones #1 and #2) and *GATA3^−/−^* (clones #1 and #2). Both *GATA3^−/−^* clones were analyzed at two different time points (early (E) and late (L)). The noradrenergic SH-SY5Y and the isogenic mesenchymal SH-EP cell lines were also included in the analysis. P-values were determined via two-tailed unpaired Welch’s t-test (**: p<0.01, ***: p<0.001).

These observations therefore demonstrated that the KO of the *PHOX2A, PHOX2B* or *GATA3* transcription factors has different consequences on the maintenance of the noradrenergic identity and that SH-SY5Y cells are able to transdifferentiate from a noradrenergic to a mesenchymal state upon *GATA3* KO.

### Spontaneous plasticity between the noradrenergic and mesenchymal states reveals the reprogramming potential of a subset of neuroblastoma cells

Our previous study has shown that most of the neuroblastoma cell lines (18 out of 25) exhibit a noradrenergic identity whereas only 3 have a mesenchymal epigenetic profile. An intermediate group of cells expressing noradrenergic TFs and mesenchymal TFs was composed of 4 samples, including the heterogeneous SK-N-SH cell line^12^. Interestingly, plasticity properties have been reported for this cell line^18^. Using single-cell transcriptomic sequencing with the 10X Genomics technology, we highlight here two cell populations in the SK-N-SH cell line. Noradrenergic cells expressed *PHOX2B* whereas *CD44*^19–21^ appeared as a specific marker of the mesenchymal population confirmed by FACS and immunofluorescence (**Figure 2A-C**). An InferCNV analysis that predict the genetic alterations at the single-cell level^22^, showed that both populations exhibited similar genetic alterations (2p, 7 and 17q gains). Only one subcluster of the mesenchymal population presented with a 1q gain (**Figure 2D**), consistently with previous data reporting distinct genetic subclones^23^. As CD44 is a cell surface marker it was further used to sort both populations. Bulk RNA-seq experiments confirmed that CD44^neg^ and CD44^pos^ sorted cells exhibited transcriptomic profiles close to noradrenergic SH-SY5Y and mesenchymal SH-EP cells, respectively (**Figure 2E**). Both sorted noradrenergic/CD44^neg^ and mesenchymal/CD44^pos^ populations were able to give rise to a heterogeneous cell population, demonstrating a spontaneous and bidirectional plasticity between the noradrenergic and mesenchymal states (**Figure 2F**). As expected from previous data obtained with the SH-SY5Y and SH-EP cell lines sub-cloned from the heterogeneous parental SK-N-SH cell line^12^, the mesenchymal/CD44^pos^ population of this cell line exhibited a higher chemo-resistance compared to the noradrenergic/CD44^neg^ one (**Figure 2G**).

**Figure 2.**
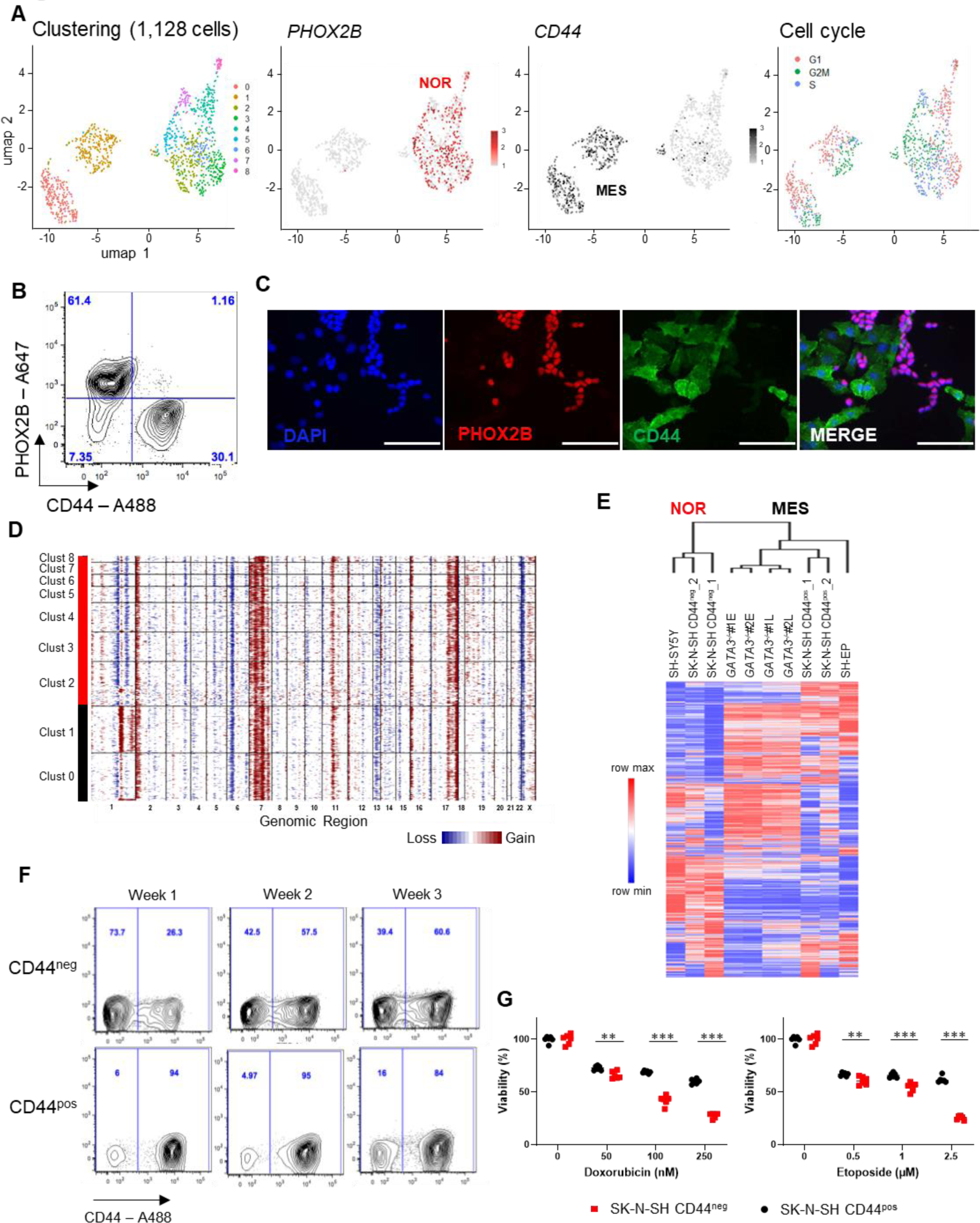
Cell plasticity in the heterogeneous neuroblastoma SK-N-SH cell line. (**A**) Single-cell RNA-seq analysis by Seurat. The umap plot shows two main clusters, *PHOX2B* and *CD44* expression distinguish noradrenergic and mesenchymal cells, each identity including cycling cells. (**B**), (**C**) FACS and immunofluorescence analyses of the SK-N-SH cell line with the PHOX2B and CD44 markers (scale bar= 50 μm). (**D**) Genetic alterations were predicted from single-cell RNAseq data using the InferCNV tool. (**E**) Unsupervised clustering using the top 10% of genes with the highest IQR (Inter-Quartile Range) indicates that CD44^neg^ and CD44^pos^ sorted cells (two replicates) exhibit a transcriptomic profile close to the noradrenergic SH-SY5Y and mesenchymal SH-EP cells, respectively. (**F**) CD44^neg^ (upper panel) and CD44^pos^ (lower panel) sorted cells are able to reconstitute a heterogeneous population of both CD44^neg^ and CD44^pos^ cells after a few weeks in culture. (**G**) Mesenchymal/CD44^pos^ sorted cells are more resistant to doxorubicin and etoposide than noradrenergic/CD44^neg^ cells. Cell viability was measured after 72 hours of chemotherapy treatments (Doxorubicin 50, 100, 250 nM and Etoposide 0.5, 1, 2.5 μM)) (mean ± sd.; n = 6 replicates). P-values were determined via two-tailed unpaired Welch’s t-test (**: p<0.01; ***: p<0.001).

In order to demonstrate that plasticity between the noradrenergic and mesenchymal states is not a property exclusively observed in the SK-N-SH sample, we generated new cell lines from a series of 10 neuroblastoma PDX models. Cells from two models could be maintained in culture for several months and frozen. One cell line had a pure noradrenergic phenotype. Strikingly, the other cell line called IC-pPDXC-63 was able to grow *in vitro* as a bi-phenotypic culture, with both adherent cells and floating neurospheres (**Figure 3A**). Bulk RNA-seq analysis confirmed that the original PDX model (IC-pPDX-63) and its derived-cell line exhibited a transcriptomic profile highly similar to the one of the patient tumor (NB1549) (**Figure 3B**). As shown by single-cell analysis, two main clusters of noradrenergic cells and mesenchymal tumor cells were observed in the IC-pPDXC-63 cell line, with a bridge of cells in-between (**Figure 3C**). Noradrenergic cells expressed *PHOX2B*, whereas the mesenchymal population expressed the cell surface CD44 marker (**Figure 3C**). The clustering separating noradrenergic and mesenchymal tumor cells is not biased by the cell cycle (**Figure 3C**). Immunofluorescence confirmed that CD44 and PHOX2B were specifically expressed by adherent cells and neurospheres, respectively (**Figure 3D**). Bulk RNA-seq analysis endorsed that CD44^pos^ FACS-sorted cells and adherent cells have a transcriptomic profile close to the mesenchymal SH-EP cells, whereas CD44^neg^ FACS-sorted cells and floating neurospheres clustered with the noradrenergic SH-SY5Y cells (**Figure 3B**). Consistently with the observations in the SK-N-SH cell line, we documented that the mesenchymal population of the IC-pPDXC-63 cell line exhibited a higher chemo-resistance compared to the noradrenergic one (**Figure 3E**). Of note, inferred genomic alterations were similar in the three populations of the IC-pPDXC-63 cell line, *i.e.*, the noradrenergic, mesenchymal and bridge cells (**Figure 3F**). We next investigated the plasticity properties of both noradrenergic and mesenchymal cells of the IC-pPDXC-63 cell line *in vitro*. After a few days in culture, both sorted noradrenergic/CD44^neg^ and mesenchymal/CD44^pos^ cells were able to reconstitute a heterogeneous cell population (**Figure 3G**), as previously documented for the SK-N-SH cell line. These observations therefore highlight the reprogramming potential between the noradrenergic and mesenchymal states of the IC-pPDXC-63 model and importantly, show that this ability does not rely on genetic heterogeneity.

**Figure 3.**
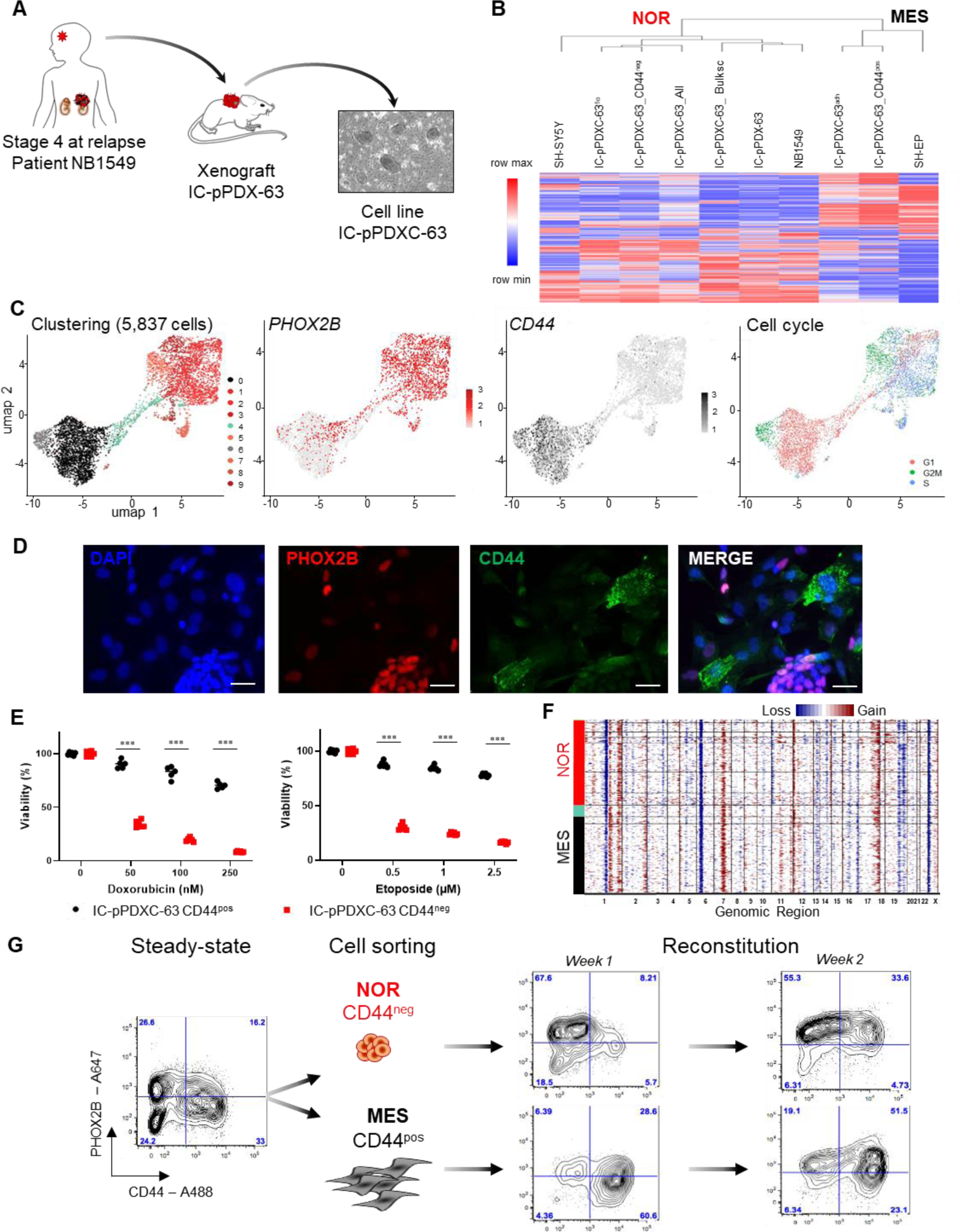
The IC-pPDXC-63 model exhibits a phenotypic plasticity between noradrenergic and mesenchymal identities. (**A**) *Ex vivo* culture of the noradrenergic IC-pPDX-63 neuroblastoma model includes floating neurospheres and adherent cells. (**B**) Unsupervised clustering using the top 10% of genes with the highest IQR (Inter-Quartile Range) shows that the transcriptomic profile of the IC-pPDX-63 model and its derived-cell line (IC-pPDXC-63) are highly similar to that of the patient’s tumor (NB1549) from which it has been generated. (**C**) Single-cell transcriptomic analyses by Seurat highlight both noradrenergic and mesenchymal clusters in the IC-pPDXC-63 cell line together with a bridge in-between. Four umap plots are shown (from left to right): the cell clustering, *PHOX2B* expression marking noradrenergic cells, *CD44* expression marking mesenchymal cells and the cell cycle. (**D**) Immunofluorescence shows the specific expression of the PHOX2B and CD44 markers by neurospheres and adherent cells, respectively (scale bar = 20 μm). (**E**) Mesenchymal (IC-pPDXC-63 CD44^pos^) cells are more resistant to chemotherapy than noradrenergic (IC-pPDXC-63 CD44^neg^) cells. Cell viability was measured with resazurin assay after 72 hours of chemotherapy treatments ((Doxorubicin 50, 100, 250 nM and Etoposide 0.5, 1, 2.5 μM)) (mean ± sd.; n = 6 replicates). P-values were determined via two-tailed unpaired Welch’s t-test (**: p<0.01; ***: p<0.001). (**F**) Inferred genomic profile of the IC-pPDXC-63 cell line obtained with InferCNV on single-cell data. (**G**) Plasticity properties of the IC-pPDXC-63 cell line. CD44^pos^ and CD44^neg^ cells were FACS sorted and put back in culture. Each sorted population reconstituted a heterogeneous cell population after several days in culture.

### Phenotypic plasticity relies on epigenetic reprogramming

To deeper characterize the epigenetic reprogramming contribution to cell plasticity, we next defined the super-enhancer landscape of our three different models (*GATA3* genetic inactivation, SK-N-SH and IC-pPDXC-63 cell lines) by ChIP-seq analyses for the H3K27ac mark. We added these samples in the principal component analysis (PCA) of neuroblastoma cell lines and hNCC lines based on their super-enhancer log scores^12^ (**Figure 4A**). For all three models, their mesenchymal counterparts, *i.e. GATA3^−/−^* clones, SK-N-SH CD44^pos^ FACS-sorted and adherent IC-pPDXC-63 cells, showed an epigenetic profile close to the group II of mesenchymal identity. Consistently, their noradrenergic counterparts, SH-SY5Y, SK-N-SH CD44^neg^ FACS-sorted and floating IC-pPDXC-63 cells were part of the noradrenergic cell line group I (**Figure 4A**). For the mesenchymal cells of the three models, a decrease of the H3K27ac signal could be quantified for the super-enhancer regions of the noradrenergic CRC such as *GATA3*, *PHOX2B* and *HAND1*. On the other hand, an increase of the H3K27ac signal could be observed on some NCC-like/mesenchymal TFs such as *RUNX1*, *FOSL1*, *NR3C1,* and *TBX18* (**Figure 4B,C**). Of note, the decreased PHOX2A and HAND2 protein expressions in the *GATA3^−/−^* clones (**Figure 1B**) contrasted with high transcript levels (**Figure 1G**) and high H3K27ac scores (**Figure 4B**) suggesting a distinct transcriptional and post-transcriptional regulation for these specific genes.

**Figure 4.**
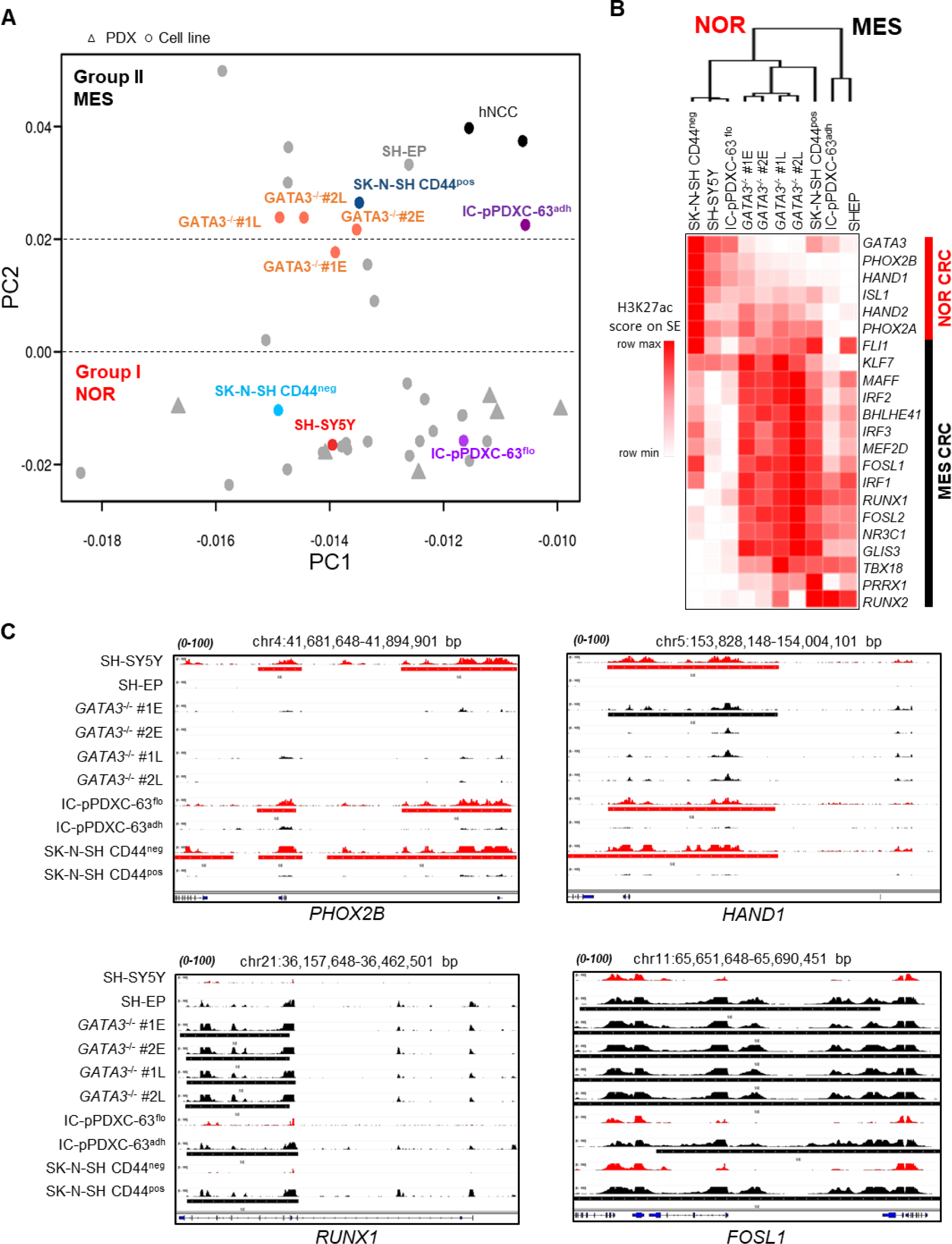
Phenotypic plasticity relies on epigenetic reprogramming. (**A**) Principal Component Analysis (PCA) based on neuroblastoma and hNCC super-enhancer log scores^12^ that discriminates the two neuroblastoma cell groups I (noradrenergic-NOR) and II (NCC-like/mesenchymal-MES) and in which were added the two *GATA3^−/−^* clones at early (E) and late (L) time points as well as the floating and adherent cells of the IC-pPDXC-63 and CD44^neg^ and CD44^pos^ sorted cells of the SK-N-SH cell lines. (**B**) Heatmap showing the H3K27ac signal on the super-enhancer regions of the TFs of the noradrenergic (NOR) and mesenchymal (MES) CRC in the two *GATA3^−/−^* at both early (E) and late (L) time points, in floating and adherent cells of the IC-pPDXC-63 cell line and in the CD44^pos^ and CD44^neg^ sorted cells of the SK-N-SH cell line, and the SH-SY5Y and SH-EP control cell lines. For the TFs associated with several super-enhancers, the signal was summarized as described in the Methods section to have one value per TF. The unsupervised hierarchical clustering based on H3K27ac signals discriminated noradrenergic and mesenchymal TFs and cell identity. (**C**) Tracks of ChIP-seq profiles for H3K27ac at *PHOX2B, HAND1, RUNX1,* and *FOSL1* super-enhancers in the SH-SY5Y, SH-EP, the 4 *GATA3^−/−^* samples (2 at early (E) and 2 at late (L) time-points), the floating and adherent IC-pPDXC-63 cells and the CD44^pos^ and CD44^neg^ sorted SK-N-SH cells.

Altogether, these results indicate that the transdifferentiation from a noradrenergic to a mesenchymal identity obtained after the genetic inactivation of *GATA3* or spontaneously from SK-N-SH and IC-pPDXC-63 cells is supported by an epigenetic reprogramming.

### Mesenchymal neuroblastoma cells are reprogrammed to a noradrenergic phenotype *in vivo*

Since noradrenergic and mesenchymal cells of our three models presented with different properties *in vitro*, we next investigated their behavior *in vivo*. We injected the noradrenergic/CD44^neg^ and mesenchymal/CD44^pos^ sorted cell populations from the two heterogeneous SK-N-SH and IC-pPDXC-63 cell lines, as well as the *GATA3^−/−^* cells, into mice. Tumors developed in all cases, indicating that the different states displayed tumorigenic potential *in vivo*. Unexpectedly, as revealed by IHC analysis, PHOX2B expression was observed in most tumor cells from the whole set of xenografts, even those obtained after engraftment of mesenchymal populations (**Figure 5A**). Bulk RNA-seq experiments confirmed that all tumors highly expressed the TFs of the noradrenergic CRC and exhibited a noradrenergic transcriptomic profile (**Figures 5B and S2**). Xenografts of the *GATA3^−/−^* clones re-expressed the noradrenergic and neuroendocrine *DBH, NET/SLC6A2, CHGA* and *CHGB* markers compared to the clones cultured *in vitro* (**Table S1**). Consistently, PHOX2B and DBH proteins were detected in the *GATA3^−/−^* cell xenografts (**Figure 5C**). Additionally, these markers were similarly expressed in xenografts obtained with the two CD44^pos^ and CD44^neg^ cell populations of the SK-N-SH and IC-pPDXC-63 cell lines (**Table S1**). Differential analyses (Fold-change >2; Bonferroni-corrected p-value <0.05) showed that xenografts obtained from CD44^pos^ and CD44^neg^ cells of both heterogeneous cell lines were highly similar, with less than 50 genes (data not shown) showing a differential expression, compared to more than 2,000 genes differentially expressed *in vitro*. Of note, some genes of the neurogenesis and extracellular matrix organization were still differentially expressed between SH-SY5Y and *GATA3^−/−^* cell xenografts, but of lower magnitude compared to the difference between the corresponding cells *in vitro*. Finally, to fully demonstrate the *in vivo* reprogramming of mesenchymal cells towards a noradrenergic identity, we determined the super-enhancer profiles of SH-SY5Y and *GATA3^−/−^* cell xenografts and additionally from the xenografts of SK-N-SH CD44^pos^ and CD44^neg^ FACS-sorted cells. The PCA clearly showed that all these tumors were part of the noradrenergic group I (**Figure 5D**). Super-enhancers marked *PHOX2B* and *HAND1* genes in xenografts of SH-SY5Y and SK-N-SH CD44^neg^ cells but also in xenografts of *GATA3^−/−^* and SK-N-SH CD44^pos^ cells (**Figure 5E**). In an *in vivo* environment, the mesenchymal cells therefore shifted back towards a noradrenergic identity indicating that the mouse microenvironment provided strong cues inducing a global epigenetic reprogramming.

**Figure 5.**
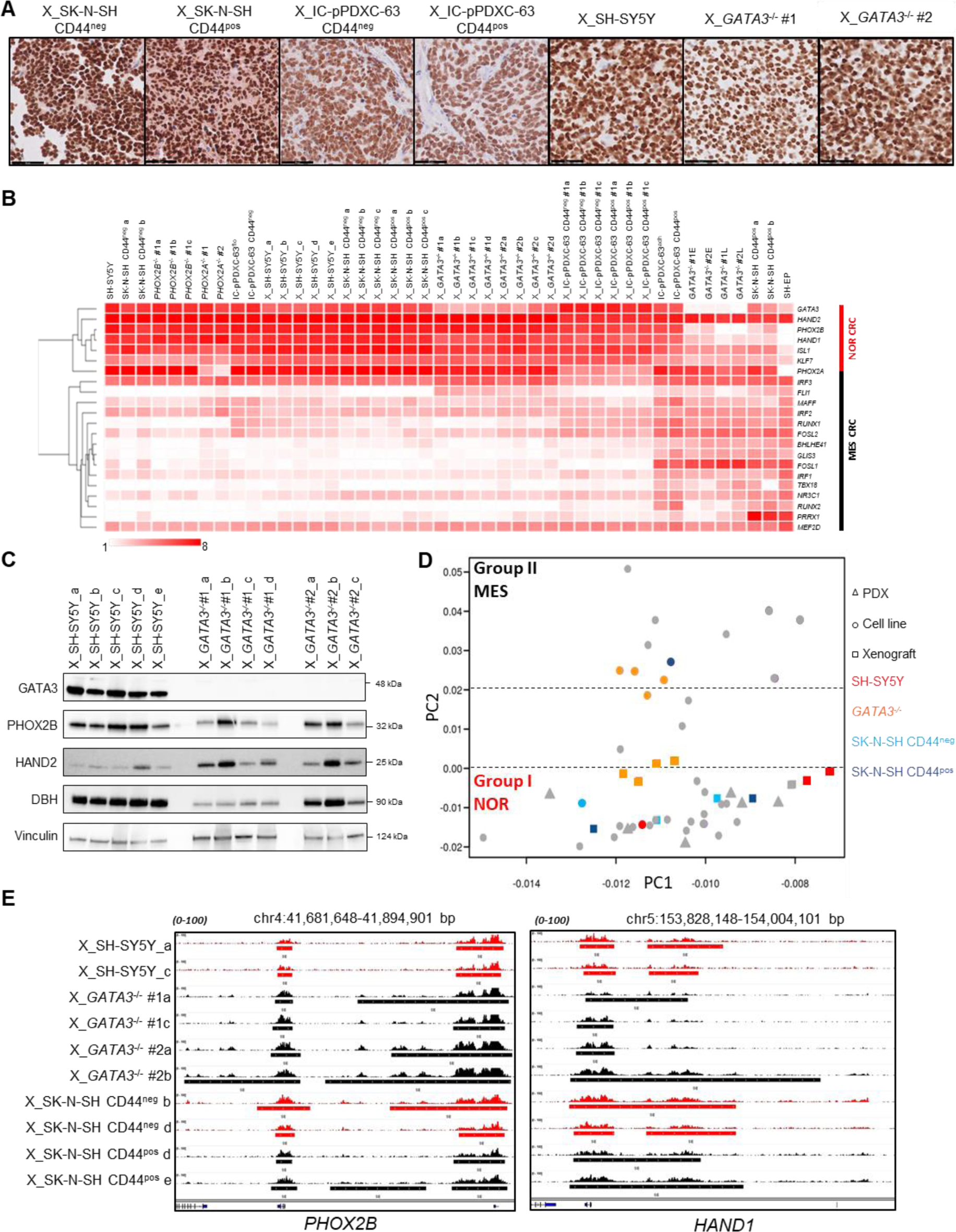
All sorted mesenchymal cells from the 3 models, SK-N-SH and IC-pPDXC-63 cell lines and *GATA3^−/−^* cells, adopt a noradrenergic identity when engrafted in mouse. (**A**) PHOX2B immunohistochemistry of one representative mouse xenograft per group (CD44^neg^ or CD44^pos^ sorted cells of SK-N-SH and IC-pPDXC-63 cell lines, SH-SY5Y control cells and *GATA3*^−/−^ clones #1 and #2). Scale bar = 50 μm. Similar results were obtained for all analyzed xenografts. (**B**) Heatmap showing the expression of the TFs associated with the noradrenergic or mesenchymal identity^12^ reveals that all mouse xenografts exhibit a noradrenergic transcriptomic profile. (**C**) Western-blot analysis of GATA3, PHOX2B, HAND2 and DBH in the SH-SY5Y (n=5) and *GATA3^−/−^* clone xenografts (*GATA3*^−/−^#1: n=4; *GATA3*^−/−^#2: n=3) with Vinculin as a loading control. (**D**) PCA as in **Figure 4A** in which were added the xenografts of the SH-SY5Y and *GATA3^−/−^* clones and the CD44^pos^ or CD44^neg^ sorted cells of the SK-N-SH cell line. (**E**) Tracks of ChIP-seq profile for H3K27ac at *PHOX2B* and *HAND1* super-enhancers in the xenografts of the following cells: SH-SY5Y, *GATA3^−/−^* clones and SK-N-SH cell populations sorted according to CD44 expression.

Altogether, our analyses highlighted the plasticity properties of neuroblastoma cells using three different models (*GATA3* genetic inactivation, SK-N-SH and IC-pPDXC-63 cell lines). This plasticity was associated with a reprogramming potential from a mesenchymal state to a noradrenergic state following *in vivo* engraftment in the mouse.

### Single-cell transcriptomic analyses reveal intra-tumor noradrenergic heterogeneity but no mesenchymal tumor cells in neuroblastoma PDX models

Since heterogeneity of cell identity has been observed in some neuroblastoma cell lines, we took advantage of the 10X Genomics technology to explore neuroblastoma intra-tumor heterogeneity using single-cell transcriptomic analyses on 14 PDX models, obtained at diagnosis, progression or at relapse (**Figure 6A, Table S2**). The study was designed to specifically identify human tumor cells (see methods). The integration of the human cells from all models (n=47,219 cells) with the Harmony tool^24^ highlighted tumor cells of noradrenergic identity (*PHOX2B*+, *HAND2+*) (**Figures 6B-C** and **S3A**). No cluster of mesenchymal tumor cells could be identified as shown by the analysis of previously published mesenchymal signatures (**Figure 6D**). The InferCNV analyses confirmed that noradrenergic tumor cells exhibited the emblematic genetic alterations of neuroblastoma such as 17q gain^1^ (**Figure S3B-C**). For several models, we could compare the InferCNV profile calculated from scRNAseq data to copy number profiles inferred from whole-exome sequencing data and confirmed their full consistency (**Figure S3B**). We next assessed the heterogeneity of the human tumor cell populations and documented that most clusters were shared by all PDX models (**Table S4**). Several populations could be defined according to the differential expression of specific genes: cycling cells marked by the expression of *TOP2A, MKI67* and *CDK1* (clusters 2 and 6), cells driven by a *MYCN/*2p amplicon signature (clusters 0-1-2-3-13), cells expressing chromaffin markers such as *CDKN1C*, *SLC18A1*/VMAT1^25,26^ and *DLK1*^27^ (clusters 9-10-11) and cells with a sympathoblast-like identity such as *TFAP2B*^28^ (cluster 7) (**Figures 6E, S3D-F** and **Table S4**).

**Figure 6.**
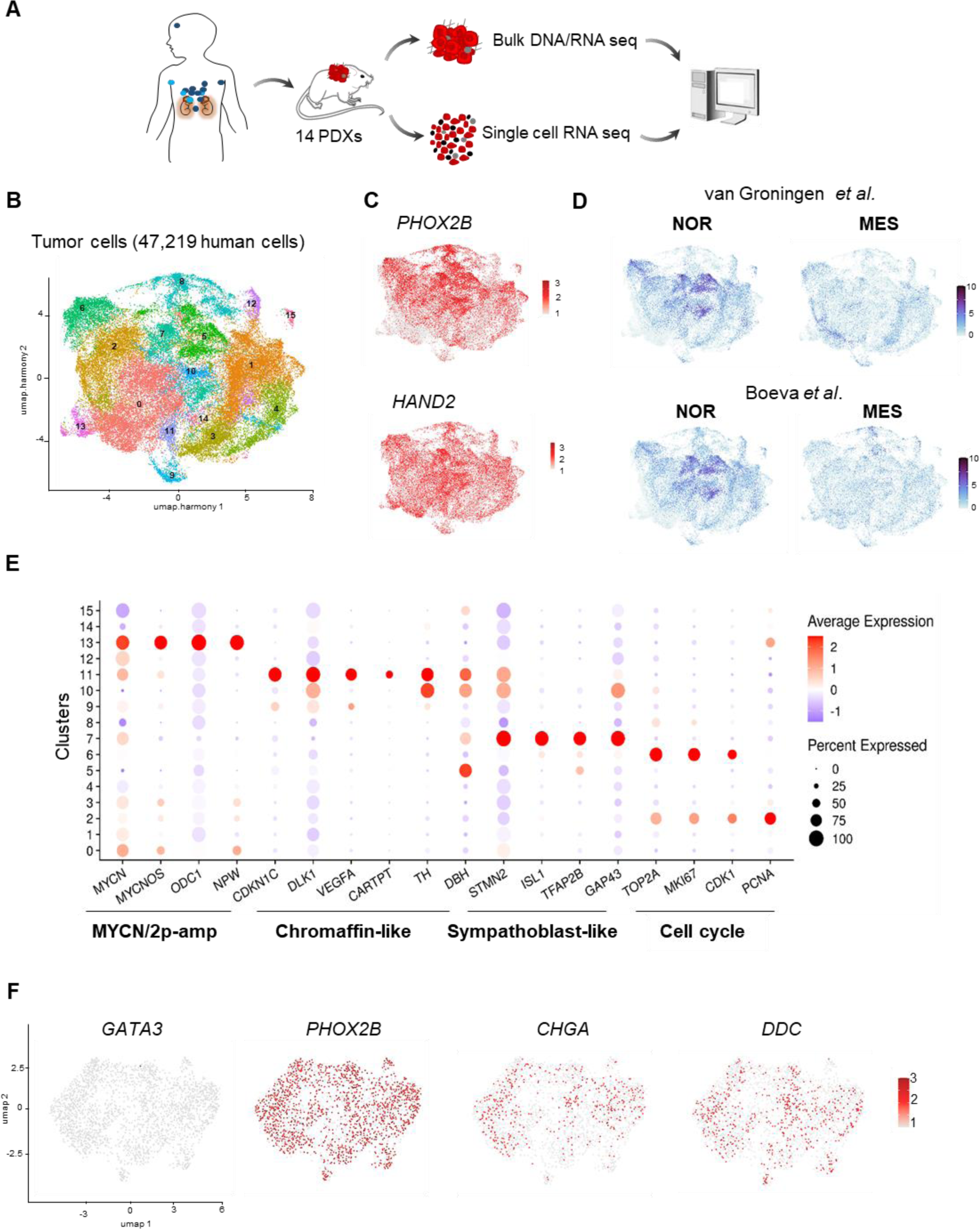
Single-cell transcriptomic analyses of 14 PDX models. (**A**) Schematic illustration of the overall procedure. scRNA-seq was performed for 14 neuroblastoma PDX models obtained either at diagnosis (light blue) or at relapse (dark blue) after tumor cell dissociation. In parallel, CNV profiles and global transcriptomes were established from the DNA and RNA of the bulk of the tumor. (**B**) Uniform manifold approximation and projection (umap) of the 47,219 human cells obtained after the integration by Harmony of the 14 PDX models. (**C**) *PHOX2B* and *HAND2* expression within the noradrenergic tumor cells. (**D**) Plots of published noradrenergic (NOR) and mesenchymal (MES) signatures^12,13^. (**E**) Dot plot graph illustrating cluster-specific gene expression. Four main populations of tumor cells could be defined. (**F**) The GR-NB7 PDX model does not express *GATA3* but exhibits a noradrenergic phenotype as shown by the plot of *PHOX2B*, *CHGA* and *DDC* expression in the scRNAseq data.

Interestingly, our single-cell transcriptomic data revealed that the GR-NB7 PDX model did not express *GATA3* but expressed *PHOX2B*, *CHGA* and *DDC* (**Figure 6F).**These data confirmed that *GATA3* was dispensable to establish a noradrenergic identity *in vivo*, in agreement with the noradrenergic shift of *GATA3*^−/−^ cells observed in mouse xenografts. Unsupervised hierarchical clustering using the 10% genes with the highest IQR on bulk RNA-seq data confirmed that all studied PDX models exhibited a transcriptomic profile corresponding to a noradrenergic identity (**Figure S3F**).

Altogether, the present analysis of 24 neuroblastoma samples identified no *bona fide* tumor mesenchymal cells, which is consistent with our aforementioned results demonstrating a strong pressure of the microenvironment towards a noradrenergic identity.

## DISCUSSION

In the present work, we have first deciphered the distinct roles of the members of the noradrenergic CRC in shaping neuroblastoma cell identity. Whereas *Phox2b* is essential for initial sympatho-adrenal cell specification from neural crest progenitors^29–31^, the KO of *PHOX2B* or *PHOX2A* did not modify the noradrenergic identity of SH-SY5Y cells. Yet, one hypothesis is functional redundancy between PHOX2A and PHOX2B. Of note, we did not succeed in obtaining double KO clones suggesting synthetic lethality for the cells. In contrast, the KO of *GATA3* in the same cells had a dramatic effect, inducing the collapse of the noradrenergic CRC. Surprisingly, whereas *GATA3* KO clones harbored mesenchymal properties *in vitro*, they reacquired a noradrenergic identity *in vivo* indicating that *GATA3* KO allows a permissive epigenetic state and the transdifferentiation towards a mesenchymal or noradrenergic state according to non-autonomous cues. Consistently with these observations, we could document a noradrenergic identity of one neuroblastoma PDX model characterized by an absence of GATA3 expression. In a previous study using siRNA and short-term transcriptomic analysis by RT-q-PCR, Durbin *et al*. reported that the knock-down of only one member of the CRC was able to induce a decrease of the expression of several members of the CRC^15^. Yet, residual expression remained for the different TFs studied and the impact of this decrease on cell identity has not been explored. Therefore, our results constitute the first evidence that TFs of a specific CRC may have distinct roles in shaping cell identity in neuroblastoma.

Transdifferentiation from the noradrenergic to the mesenchymal identity has been shown previously *in vitro* following overexpression of the PRRX1 TF^13^. The same team also recently pointed out a role of the NOTCH signaling pathway in this process^32^. Of note, xenografts of SH-SY5Y cells overexpressing NOTCH3 intracellular domain exhibited a mesenchymal identity. In this model, the permanent production of intra-cellular NOTCH3 likely activates an endogenous feed-forward loop between NOTCH receptors and ligands *in vivo.* Although we observed expression of several members of this pathway (*NOTCH2*, *MAML2*, *HES1*) in our mesenchymal tumor cells *in vitro*, in our series of PDX models, only expression of *MAML2* could be detected in some tumor cells; yet these cells were not associated with a specific cluster and very spare expression was observed for *NOTCH2* and *HES1* in the whole population of tumor cells.

We document here the heterogeneity and spontaneous reprogramming potential of cells of IC-PDXC-63 and SK-N-SH models from a noradrenergic to a mesenchymal identity and conversely. Single-cell transcriptomic analysis performed on these models first identified CD44 as a surface marker specific of the mesenchymal identity, further allowing the use of this marker to sort each population and analyze their respective proportions by FACS. We show that the spontaneous and bi-directional plasticity observed *in vitro* relies on a profound epigenetic reprogramming, as revealed by the analysis of the super-enhancer landscape of the various cell populations. Very strikingly, we demonstrate that mesenchymal tumor cells from three different models exhibiting plasticity revert their identity towards a noradrenergic state when engrafted in mice. These results highlight that neuroblastoma cell phenotype is strongly influenced by the *in vivo* microenvironment which provides a powerful pressure towards a noradrenergic state. This conclusion is reinforced by the observation that mesenchymal tumor cells were not identified by single-cell transcriptomic analyses in our series of 14 neuroblastoma PDX models, obtained either at diagnosis or at relapse and presenting various genetic alterations. Altogether, our data obtained on a variety of cellular models and PDXs provide a biological explanation for the absence of mesenchymal tumor cells *in vivo*. This observation is also in line with the very recent study performed by Dong and colleagues that identified normal, but not tumor, mesenchymal cells in adrenal neuroblastoma tumors by single-cell RNA sequencing^33^. Interactions between tumor cells and several cell populations of the microenvironment may influence the tumor cell phenotype and play a role in tumor progression, as previously demonstrated for tumor-associated inflammatory cells^34^. Future studies should decipher the cues of the microenvironment and their associated pathways that converge to regulate cell plasticity during tumor progression.

Interestingly, some evidences suggested that treatments against neuroblastoma may impact cell identity. Indeed, previous analyses of neuroblastoma cells selected to be resistant to cisplatin^35^ or ALK inhibitors^36^ *in vitro* have reported that noradrenergic cells may acquire mesenchymal properties. Cells exhibiting such a potential may be transiently induced upon chemotherapy treatment and revert their state towards a noradrenergic identity when the treatment pressure decreases, the mesenchymal cells being the drug-resistant reservoirs for noradrenergic cells. Further experiments will allow better defining their contribution to therapeutic resistance and relapse in high-risk neuroblastomas.

The cell of origin in neuroblastoma has been for long a matter of debate^37–39^. Some models argue that this tumor arises from the transformation of NCCs while other models suggest that they develop from more engaged sympatho-adrenal progenitors, able to generate both sympathetic neurons and neuroendocrine cells of the adrenal. Recently, this hierarchical dogma of normal differentiation has been questioned with the identification of a population of Schwann cell precursors (SCPs) as the main reservoir of adrenal chromaffin cells in the mouse^26^. In their recent paper, Dong et al. who analyzed human fetal adrenal gland at the single-cell level, in addition to adrenal neuroblastoma tumors, concluded that malignant cells had a predominant chromaffin-cell-like phenotype^33^. We nevertheless disagree with their interpretation of cell phenotypes. We strongly believe that *CARTPT* and *INSM1,* used by Dong et al. to define sympathoblast identity, rather witness a chromaffin-like phenotype, since they are co-expressed with *DLK1*, *CDKN1C*, *CHGA*/*CHGB*, *TH*, *DBH* and *SLC18A1* in our data ^25–27,40–44^. Integration of our single-cell RNA-seq data from 14 PDX models revealed two distinct cell populations expressing either markers of chromaffin cells or markers of sympathoblasts. Of note, SCP markers such as *SOX10, S100B, PLP1* and *ERBB3*^26^ were not detected in our cohort of 14 neuroblastoma cases. Nevertheless, it remains difficult to infer which cell type is the one targeted by neoplastic transformation. Indeed, it cannot be excluded that transitions may occur between chromaffin cells and sympathoblasts during development and/or that markers of a specific precursor may be lost during cell transformation.

Altogether, our data obtained on several cellular models demonstrate that a subset of neuroblastoma cells exhibits a reprogramming potential between a noradrenergic and a mesenchymal identity and that both intrinsic properties and exogenous signals of the microenvironment dictate this identity. A better understanding of the molecular factors that control phenotypic plasticity will represent a key step in the design of more efficacious therapies that aim at improving the outcome of neuroblastoma patients with high-risk disease.

## ACKNOWLEDGEMENTS

This work was supported by grants from Institut Curie, Inserm, the Ligue Nationale Contre le Cancer (Equipe labellisée), the Institut National du Cancer (PLBIO18-273) and by the following associations: Association Hubert Gouin-Enfance et Cancer, Les Bagouz à Manon, les amis de Claire, Courir pour Mathieu, Dans les pas du Géant, Olivier Chape. The Mappyacts protocol is supported by the Institut National de Cancer (INCa) through the PHRC “INCa-DGOS_8519” MERRI, the Fondation ARC, the Association Imagine for Margo, the Société Française de lutte contre les Cancers et les leucémies de l’Enfant et l’adolescent (SFCE), Fédération Enfants & Santé, the associations AREMIG and Thibault BRIET. The Micchado protocol is supported by PRT-K, Association Imagine for Margo, Kickcancer, Hubert Gouin-Enfance et Cancer, Fédération Enfants & Santé, and funding support by BMS and Roche. High-throughput sequencing has been performed by the ICGex NGS platform of the Institut Curie supported by the grants ANR-10-EQPX-03 (Equipex) and ANR-10-INBS-09-08 (France Génomique Consortium) from the Agence Nationale de la Recherche (“Investissements d’Avenir" program), by ITMO Cancer Aviesan (“Equipement pour la recherche en Cancérologie” program) and by the SiRIC-Curie program -SiRIC Grant “INCa-DGOS- 4654”. G.S. is supported by the Annenberg foundation. H.R. is supported by the Wilhelm-Sander-Stiftung. We are grateful to the animal facilities team, the Experimental Pathology Department, the Plateforme Génomique and the Plateforme Cytométrie of Institut Curie, U900 colleagues for help with alignment of NGS data and Julien Masliah-Planchon for help with genomic analyses. We thank Mélissa Saichi and Divya Sahu for preliminary analyses of single-cell RNA-seq and ChIP-seq data.

## AUTHOR CONTRIBUTIONS

Conceptualization: C.T., V.B., O.D. and I.J.-L.; Methodology: C.T., A.P., S. Z., N.G., I. J., A.M.C., B.G. and D.S.; Formal Analysis: C.T., A.K., A.C., S.G.-L., V.B.; Investigation: C.T., A.P., S.D., C.L.-B., C.P.-E., A.C., A.G., N.G. and H.R.; Resources: E.L., G.P., H.B., A.G., P.F., L.G. B., V.R., S.B., A. B., J.B and G.S.; Writing – Original Draft: C.T., A.P. and I.J.-L.; Writing – Review and Editing: C.T., A.P., C.L.-B., H.R., O.D. and I.J.-L.; Visualization: C.T., A.P., A.K., C.L.-B., C.P.-E.; Supervision: C.T. and I.J.-L.; Project Administration: I.J.-L.; Funding Acquisition: I.J.-L, V. B. and O.D.

## MATERIAL AND METHODS

### RESOURCE AVAILABILITY

#### Lead Contact

Further information and request for resources should be directed to and will be fulfilled by the Lead Contact, Isabelle Janoueix-Lerosey.

#### Material availability

Availability of the IC-pPDXC-63 model generated in this study is subjected to a Material Transfer Agreement.

#### Data and code availability

ChIP-seq (RRID:SCR_001237) data of cell lines and xenografts (34 samples) are available in Gene Expression Omnibus (GEO, RRID:SCR_005012).

All single-cell RNA-seq from PDXs, all RNA-seq and the ChIP-seq data on the IC-pPDXC-63 cell line are available in European Genome-Phenome Archive (EGA) under the accession number EGAS00001004781 (ongoing submission).

### EXPERIMENTAL MODELS

#### Neuroblastoma cell lines

The SK-N-SH (Cat# HTB-11, RRID:CVCL_0531) and SH-SY5Y (Cat# CRL-2266, RRID:CVCL_0019) cell lines have been obtained from the ATCC. The SH-EP cell line has been kindly provided by M. Schwab. Cell line authentication was done by STR profiling with PowerPlex® 16 HS System from Promega. The IC-pPDXC-63 cell line was derived from the IC-pPDX-63 model. *GATA3*^−/−^ clones were genetically modified from the SH-SY5Y cell line. Cells were grown at 37°C with 5% CO_2_ in a humidified atmosphere in DMEM/HIGH glucose (Cat# SH30022.01, GE Healthcare) for SK-N-SH, SH-SY5Y and *GATA3*^−/−^ clones and in RPMI-1640 (Cat# SH30027.01, GE Healthcare) for IC-pPDXC-63 and SH-EP cell lines, with 10% FBS (Cat# SV30160.03, GE Healthcare). Cells were monthly checked by qPCR (Venor® GeM qEP 11-9250, Minerva biolabs®) for the absence of mycoplasma.

#### Mouse xenograft experiments

For each cell line, 5 million cells were injected subcutaneously in the flanks of 8 week-old Nude or NSG mice with a ratio of 50/50 standard medium (DMEM: Cat# SH30022.01, GE Healthcare) and BD Matrigel™ (Cat# 356234, BD Biosciences). Tumor volume was measured every 2 or 3 days with a caliper. Mice were sacrificed when the tumor reached a volume of 2,000 mm^3^ calculated as V = (a/2) * b * ((a+b)/2), a and b being the largest and smallest diameters, respectively.

*In vivo* experiments for this study were performed in accordance with the recommendations of the European Community (2010/63/UE) for the care and use of laboratory animals. Experimental procedures were specifically approved by the ethics committee of the Institut Curie CEEA-IC #118 (Authorization APAFIS#11206-2017090816044613-v2 given by National Authority) in compliance with the international guidelines.

#### Patient-derived Xenografts (PDX models)

Written informed consents for the establishment of PDXs were obtained for all patients from parents or guardians. GR-NB4 (previously named MAP-GR-A99-NB-1^12^), GR-NB5 (previously named MAP-GR-B25-NB-1^12^), GR-NB7 and GR-NB10 have been provided by Birgit Geoerger (Gustave Roussy, Villejuif, France). IC-pPDX-63, IC-pPDX-75, IC-pPDX-109 and IC-pPDX-112 have been developed at Institut Curie. HSJD-NB-003, HSJD-NB-004, HSJD-NB-005, HSJD-NB-007, HSJD-NB-009 and HSJD-NB-011 PDX models have been provided by Angel Carcaboso (Institut de Recerca San Joan de Déu, Barcelona, Spain). These models have been generated from patients under an Institutional Review Board-approved protocol or within a clinical trial, respectively (**Table S2**).

### METHOD DETAILS

#### CRISPR-Cas9 KO strategy

The guide RNAs used to specifically target the *PHOX2A*, *PHOX2B* and *GATA3* genes were selected from the CRISPOR website (crispor.tefor.net) for their high predicted efficiency and specificity: CAATTCGTACGACTCGTGCG and CTTGGAATCGTCGTCCTCGG targeting *PHOX2A*, CCCAGCCATACAGGACTCGT and AAACTCTTCACGGACCACGG targeting *PHOX2B*, GTACTGCGCCGCGTCCATGT and GAGCTGTACTCGGGCACGTA targeting *GATA3*. *PHOX2A, PHOX2B* and *GATA3* KO were performed by inducing a large deletion between exons 1 and 3, exons 1 and 2 or exons 2 and 3, respectively (**Figure S1A**). To screen the KO clones, 3 pairs of primers were used per gene: surrounding the deleted region (*PHOX2A*: 5’-CCGATGGACTACTCCTACCTC-3’, 5’-GCCGGCAGCTAGAAGAGATT-3’, *PHOX2B*:5’-GTTGGACAGCTCAGTTCCC-3’, 5’-CCCTAGGTCCTTCTCACTCG-3’,*GATA3*:5’-GCAGAATTGCAGAGTCGTCG-3’,5’-AAGAGCTGGCTCCTACCTGT-3’), at the 1^st^ cut site (*PHOX2A*: 5’-GGCCGATGGACTACTCCTACCT-3’, 5’-GGGGGACAGTCGCATTCAC-3’, *PHOX2B*: 5’-CAGCAATAAGACCAACCGCT-3’, 5’-GGTTCGGGTGTGACTAGGAT-3’, *GATA3*: 5’-TTGCTAAACGACCCCTCCA-3’, 5’-AAATGAACCAGGAACGGCAG-3’) and at the 2^nd^ cut site (*PHOX2A*: 5’-AGCTTTGAAAACCCGGAGCC-3’, 5’-CGGCTGCCAAGCCTTAAGTA-3’, *PHOX2B*: 5’-TCTCAAGTCCGTCACATCGC-3’, 5’-ATTTCTGATCGGCCATGGGG-3’, *GATA3*: 5’-TGCGAGGTAGAGATTCCCCA-3’, 5’-GCTAGGATGGGAGGACATGC-3’). The guide RNAs, crRNA and the recombinant Cas9 protein were purchased from Integrated DNA Technologies. Their reconstitution, assembly and delivery to the cells were performed following manufacturer’s instructions. KO efficiency was determined 2 days after cell transfection and clones were generated by plating 200 cells in 10 cm culture dishes. KO clones were verified by Sanger sequencing with the BigDye Terminator V1.1 Cycle Sequencing Kit (130-098-462, Thermo Fisher).

#### Phalloidin staining

100,000 cells for SH-SY5Y, *PHOX2B*^−/−^ and *PHOX2A*^−/−^ clones, 50,000 cells for *GATA3*^−/−^ clones were plated in a 4-well Lab-Tek chamber (Cat# 177399PK, Thermo Fisher) 24 hours before immunostaining. Cells were fixed with 4% PFA buffer, permeabilized with a 0.2% triton solution and blocked in a 1% BSA 0.1% triton solution. Phalloidin–Tetramethylrhodamine B isothiocyanate (Sigma Aldrich Cat# P1951, RRID:AB_2315148) was used at 1:100 to stain actin filaments. DAPI (Cat# 62248, Thermo Fisher) was diluted at 1: 1,000 in ProLong™ Gold (Cat# P36930, Thermo Fisher).

#### Immunoblotting

Proteins were extracted using a RIPA buffer (NaCl 150m M, Tris 50 mM pH=7.5, EDTA 1 mM, SDS 0.1%, deoxycholic acid 0.25%, IGEPAL 1%, PMSF 1 mM) supplemented with protease inhibitor cocktail tablets (Cat# 11836145001, Roche). 30 to 40 μg of proteins were used for Western blot analysis. The antibodies targeting the PHOX2B N-terminal part (Santa Cruz Biotechnology Cat# sc-376997, RRID:AB_2813765), PHOX2A (Santa Cruz Biotechnology Cat# sc-81978, RRID:AB_1127226) and DBH (Santa Cruz Biotechnology Cat# sc-15318) were used at 1:500. Anti-GATA3 (Cell Signaling Technology Cat#5852) and HAND2 (Abcam Cat# ab200040) were used at 1: 1,000 and anti-Vinculin (Abcam Cat# ab129002, RRID:AB_11144129) was used at 1: 10,000.

#### Proliferation assays

To evaluate the proliferation rate of the *PHOX2B^−/−^* and *GATA3^−/−^* clones compared to the parental SH-SY5Y cell line, 10,000 cells were plated in a 24-well plate in 6 replicates. Cells were then counted at day 3, 7 and 10, using a Vi-cell XR Viability Analyzer (Beckman Coulter).

#### Invasion and migration assays

Invasion assays were performed in 96-well low adherent plates (Corning) with 2,000 cells per well for the SH-EP cell line (n=12), and 10,000 cells per well for the SH-SY5Y cell line (n=12) and *GATA3^−/−^* clones (n=18). After 4 days of culture, spheroids were embedded in Corning® Collagen I (cat# 354249, Corning) and cultured in normal conditions. Invasive cells were observed 72 hours after spheroid inclusion and the area of migration was quantified using the ImageJ software (RRID:SCR_003070).

Transwell experiments were performed using 12-well 8 μm culture inserts (Dutscher). 50,000 cells were used per condition. After 24 hours of incubation, cells that have migrated through the insert membrane were fixed and stained with crystal violet (Sigma Aldrich). In each replicate, migrated cells were quantified using the mean of 6 different images, 6 replicates were performed per cell line.

#### Immunofluorescence

100,000 cells for SK-N-SH and IC-pPDXC-63 cell lines were plated in a 4-well Lab-Tek chamber (Cat# 177399PK, Thermo Fisher) 48 hours before immunostaining. Cells were fixed with 4% PFA buffer, permeabilized with a 0.2% triton solution and blocked in a 1% BSA 0.1% triton solution and incubated with anti-PHOX2B (Santa Cruz Biotechnology Cat# sc-376997, RRID:AB_2813765) and anti-CD44 (Cat# 15675-1-AP, Proteintech) at 1:100. Secondary antibodies were Cy5-Anti-Mouse (Cat# 715-175-151, Jackson ImmunoResearch Labs / 1:100, RRID:AB_2340820) and Cy3-Anti-Rabbit (Cat# 711-165-152, Jackson ImmunoResearch Labs / 1:100, RRID:AB_2307443) and DAPI (Cat# 62248, Thermo Fisher) was diluted at 1: 1,000 in ProLong™ Gold (Cat# P36930, Thermo Fisher).

#### Chemotherapy treatments

SH-SY5Y, SH-EP or *GATA3*^−/−^ clones were plated in a 24-well plate 24 hours before the chemotherapy treatments. Seeding densities for each cell lines were optimized to reach 80% confluence in the untreated cells. Cells were treated with 3 different doses of the conventional chemotherapy for 2 days and cell viability was assessed using a Vi-cell XR Viability Analyzer (Beckman Coulter); 6 replicates were performed for each cell line and each dose.

IC-pPDXC-63 and SK-N-SH cell lines were plated in 96-well plates 24 hours before the addition of doxorubicin or etoposide. Seeding densities for each cell line were optimized to reach 80% confluence in the untreated cells. Cells were treated with chemotherapeutic agents for 72 hours. Cell viability was then measured using the Resazurin reagent (Sigma-Aldrich).

#### RNA-sequencing and analyses

RNAs were extracted from frozen tumors by mechanical crushing followed by TRIzol® reagent (Cat#15596018, Invitrogen) and purified with the NucleoSpin RNA kit (Cat# 740955.50, Macherey-Nagel). For the bulk scRNA-seq samples and the cell lines, extraction and purification were done directly using this NucleoSpin RNA kit. RNA quality was assessed with a Bioanalyzer instrument and RNAs with an RNA Integrity Number above 7 were processed for sequencing as previously described^12^. RNA sequencing libraries were prepared from 500 ng to 1 μg of total RNA using the Illumina TruSeq Stranded mRNA Library preparation kit according to manufacturer recommendation. For the xenografts of IC-pPDXC-63 *in vivo* sample, mRNA Library preparation was done with TruSeq RNA Exome from Illumina. 100 bp paired-end sequencing was performed with the Illumina NovaSeq 6000 instrument (pair-ended, 100 nt).

Reads were aligned to the human reference genome hg38/GRCh38 using STAR 2.6.1a_08-27 (RRID:SCR_015899, https://github.com/alexdobin/STAR) with the following options: -outFilterMismatchNoverLmax 0.04 --alignIntronMin 20 – lignIntronMax 1000000 – outFilterMultimapNmax 20. Gene expression values (FPKM=fragments per kilobase per million reads) were computed by Cufflinks v2.2.146 (RRID:SCR_014597, http://cole-trapnell-lab.github.io/cufflinks/) and further normalization between samples was done using quantile normalization (R/Bioconductor package LIMMA (RRID:SCR_010943)).

#### PHOX2B immunohistochemistry

Tumors were fixed in a 4% formol buffer (VWR) during 24 hours, embedded in paraffin and cut in 4 μm slices. For PHOX2B immunohistochemistry, the REAL™ EnVision™ Detection System (Cat# K406511-2, Agilent Technologies) was used and the antibody against PHOX2B (Cat# sc-376997, Santa Cruz) was diluted at 1:200. Xenograft slices were also colored with a hematoxylin solution.

#### ChIP-seq and analyses

H3K27ac chromatin immunoprecipitation (ChIP) experiments were performed as previously described^12^ using the iDeal ChIP-seq kit for histones (Cat# C01010171, Diagenode) and the H3K27ac rabbit polyclonal antibody (Abcam Cat# ab4729, RRID:AB_2118291). Illumina sequencing libraries were prepared from the ChIP and input DNA and sequenced on the Illumina NovaSeq 6000 instrument (single reads, 100 nt). ChIP-seq reads were mapped to the human reference genome hg19/GRCh37 using Bowtie2 v2.1.0 (http://bowtie-bio.sourceforge.net/bowtie2/index.shtml) and further analyzed with HMCan v1.40 (RRID:SCR_010858)^45^. Super-enhancers were called with LILY software as previously described^12^. LILY was also used to normalize HMCan density profiles between samples. The H3K27ac signal on super-enhancers shown in the heatmap was computed as the sum of normalized H3K27ac densities divided by the length of the super-enhancers. For TF genes associated with several super-enhancers, the signal of the associated super-enhancers is summed and divided by their total length. PCA was computed as described previously^12^.

#### Flow cytometry and sorting

Flow cytometry analysis was performed with the BD™ LSRII cytometer. Cells were detached with TrypLE™ Express Enzyme (Cat# 12604013, Gibco), suspended in PBS and permeabilized with the IntraPrep kit (Cat# A07803, BeckmanCoulter). The cell suspension was incubated with PHOX2B [Clone B-11]-AlexaFluor® 647 (Cat# SC-376997 AF647, Santa Cruz) and CD44-FITC (Cat# 103005, Biolegend, RRID:AB_312956) antibodies during 40 min at 4°C.

Flow cytometry sorting was performed with the S3e™ cell sorter (Bio Rad). Cells were detached with TrypLE™ Express Enzyme (Cat# 12604013, Gibco), suspended in PBS and incubated with CD44-FITC antibody 30 min at 4°C in dark. Cells positive and negative for CD44 staining were sorted.

The first gating based on FSC/SSC represents 60% for IC-pPDXC-63 and 75% for SK-N-SH. Doublet cells are eliminated by gating on SSC-W / SSC-H followed by FSC-W / FSC- H. The second gating based on DAPI negative staining eliminates dead cells. The boundaries between positive staining and negative staining are always more than 1 Log of fluorescence intensity. A control tube without staining is always analyzed to determine auto-fluorescence.

#### Tumor dissociation into single-cell suspension

PDX tumors were cut with scalpels in small fragments. Enzymatic dissociation was realized in CO_2_ independent medium (GIBCO) containing 150 μg/mL Liberase™ TL Research Grade (Cat# 5401020001, Merk) and 150 μg/mL DNase (DN25, Sigma Aldrich), for 30 min at 37°C with 400 rpm agitation. Cell suspension was then filtered using 70 μm cell strainer (Cat# 130-098-462, Miltenyi Biotec). The cell suspension was washed twice with PBS. Viability was measured using Vi-cell XR Viability Analyzer (Beckman Coulter). For some PDXs, the Mouse Cell Depletion KIT was used following the manufacturer’s instructions (Cat# 130-104-694, Miltenyi Biotec).

#### Single-cell RNA-sequencing experiments and preprocessing of data

Single-cell RNA-seq was performed with the 10x Genomics Chromium Single Cell 3’ Kit (v3) according to the standard protocol. Libraries were sequenced on an Illumina HiSeq2500 or NovaSeq 6000 sequencing platform.

CellRanger version 3.1.0 (10x Genomics, https://support.10xgenomics.com/) was used to demultiplex, align and generate UMI count tables from sequencing reads. Two reference genomes were used to align reads:

- The human reference genome (hg38/GRCh38) for *in vitro* cell lines.
- A human-mouse reference genome (GRCh38-mm10) for the 14 neuroblastoma PDX models. In this scenario, we identified the mouse and human cells after inspection of the percentage of coverage from GRCh38 genome. We labeled cells as either human (at least 80%), murine (less than 30%) or human-murine doublets (between 30-80%). Only human and human-murine doublet cells were selected and coverage plus gene information of only GRCh38 genome were retained for downstream analysis.

Summary of analyses are shown in **Table S3**. Of note, we performed two technical replicates for three models: HSJD-NB-005, IC-pPDX-63 and IC-pPDX-75 (* in Table S3) to assess reproducibility at distinct passages in mice (data not shown).

#### Doublet detection

Scrublet^46^ v0.2.1 was used to detect potential doublets using default parameters (expected_doublet_rate=0.06). Cells marked as doublets were removed from subsequent analysis (results in **Table S3**). Doublet detection was only feasible on samples aligned to GRCh38 genome.

#### Quality control of single-cell data

First, all ribosomal genes (defined as *RLP/RPS* genes) were removed from the raw expression matrices. Then, coverage thresholds were set for each sample individually; an upper threshold was set to remove outlier cells with coverage greater than the 99th percentile, and a lower threshold was set to remove low quality cells with coverage inferior to the 1st percentile, except for the IC-pPDXC-63 cell line for which the limits were 1000 UMI and 500 genes detected per cell. To avoid cells with low number of genes, the same lower threshold was applied on the number of genes thus defining a minimum number of genes required. Finally, cells with more than 20% of reads mapping mitochondrial genes were removed.

#### Normalization of single-cell data

Raw UMI counts were normalized using the “SCTransform” function of Seurat^47,48^ v3.1.5 (RRID:SCR_007322). Regressed variables included cell coverage, number of features, and the percentage of UMI from mitochondrial genes.

#### Dimensionality reduction and cluster identification

Normalized count data was subjected to dimensionality reduction keeping the first 30 principal components. Uniform Manifold Approximation and Projections (umap) embeddings were calculated using these PCs as input and cells were clustered using the “FindClusters” function of Seurat.

#### Cell type annotation

Marker genes that define cell clusters were identified after differential expression analysis using Seurat’s “FindAllMarkers” function. Clusters were annotated by comparing their top marker genes to canonical cell type markers from the literature.

#### Generation of single-cell signature scores

To plot the expression of gene signatures in single cells, we used the “AddModuleScore” function from Seurat R package with 100 genes in the control gene set. Expression scale was binned into 10 bins when a gene signature is plotted and into 3 bins (1= low, 2= median, 3= high) when a single gene is plotted.

#### scRNA-seq data integrations

Harmony^24^ (https://github.com/immunogenomics/harmony) was used to integrated the 14 PDX models. Downstream analysis was carried out as described in ‘’Dimensionality reduction and cluster identification”. Of note, when several single-cell transcriptomes are available for the same model, only one was used for the integration to avoid over-representation.

#### Cell cycle analysis

We scored single cells based on expression of G2/M and S phase markers using Seurat’s “CellCycleScoring” function.

#### Copy number analysis in single cells

Copy number variations at the single cell level were called with R package InferCNV v1.2.1 ^22^ (https://github.com/broadinstitute/inferCNV) using default parameters. Normal cells from the microenvironment were used as reference cells. Cells with fewer than 1000 UMI were excluded and monocytes from publicly available single cell RNA sequencing of healthy human PBMCs^49,50^ (GEO: GSE115189) were used as reference cells (n=376).

### QUANTIFICATION AND STATISTICAL ANALYSES

Statistical tests were performed using GraphPad Prism 8 (RRID:SCR_002798). Significance values are described in the figure legends. P-values were determined via two-tailed unpaired Welch’s t-test (**:p<0.01; ***:p<0.001).

## SUPPLEMENTAL INFORMATION

**Figure S1:**
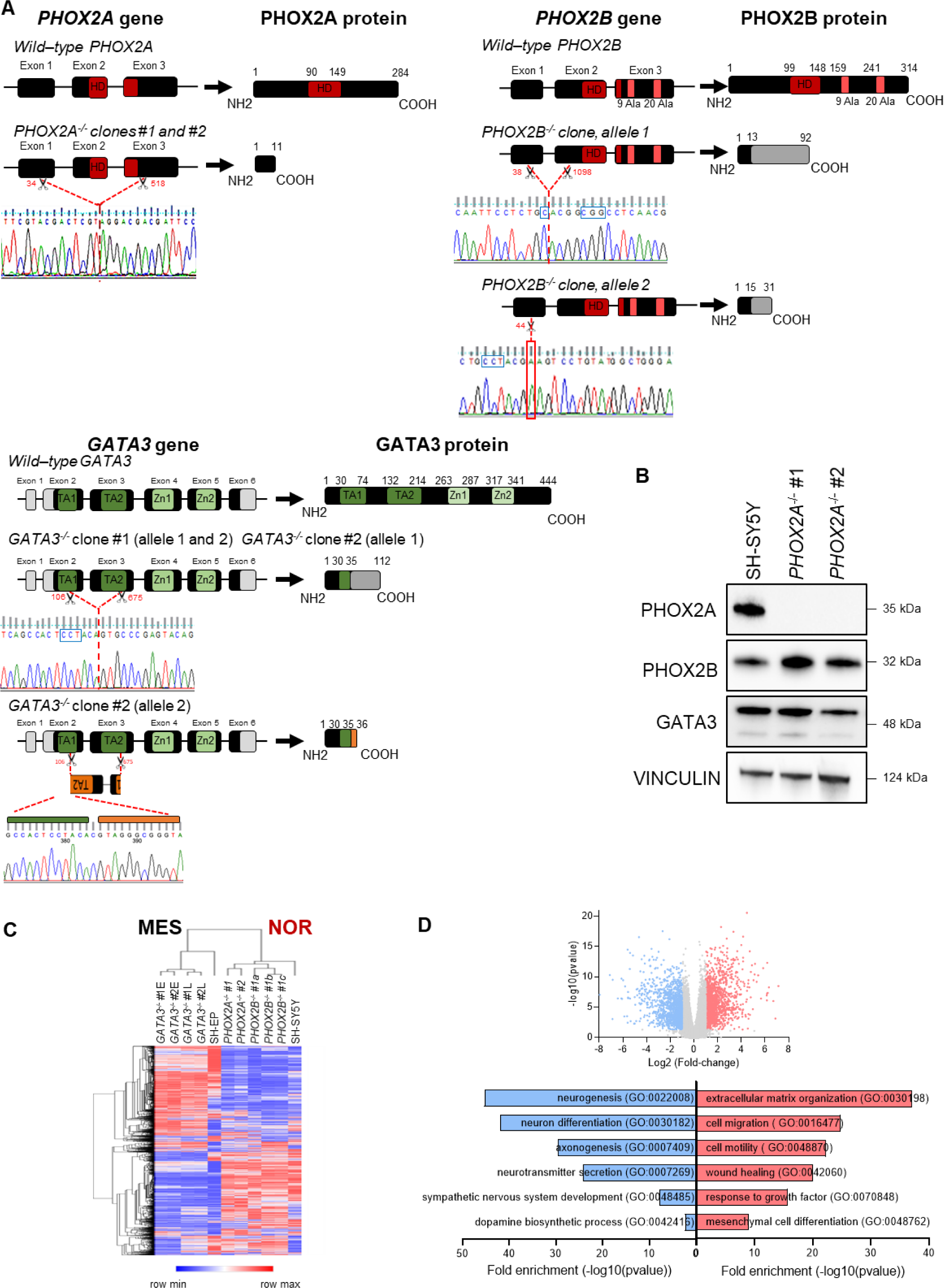
Knock-out of *PHOX2A, PHOX2B* and *GATA3* genes by CRISPR-Cas9. (**A**) Two guide RNAs were chosen per gene to induce large deletions (see Methods). The PAM motifs are surrounded in blue when still presents in the modified sequence. Both *PHOX2A*^−/−^ clones are homozygous and present with a large deletion between the 1^st^ and 3^rd^ exons, following Cas9-induced breaks at the expected positions. This results in a frameshift and subsequently in a truncated protein that has lost all its functional domains, containing only the 11 first amino acids of the normal PHOX2A protein. The *PHOX2B*^−/−^ clone is heterozygous. In the first allele, the Cas9 induced one break in exon 1 in the PAM motif instead of 3 bp upstream and one break in exon 2, 4 bp upstream of the PAM (instead of 3 bp upstream). This results in a frameshift and subsequently in a truncated protein, in which only the first 13 amino acids of the normal PHOX2B protein are conserved. In the second allele, the Cas9 induced only one cleave in the first exon, 3 bp upstream of the PAM, as expected and an “A” insertion was observed. This results in a frameshift and subsequently in a truncated protein, in which only the first 15 amino acids of the normal PHOX2B protein are conserved. *GATA3*^−/−^ clone #1 is homozygous and presents with a large deletion between the 2^nd^ and 3^rd^ exons, following Cas9-induced breaks that occurred at the expected positions. This leads to a truncated protein of 112 amino acids that has lost all its functional domains and contains only the 35 first amino acids of the normal GATA3 protein. *GATA3*^−/−^ clone #2 is heterozygous. One allele is similar to those of *GATA3*^−/−^ clone #1. For the second allele, the break occurred at the expected position but the *GATA3* sequence between these two cleavage sites was reintegrated in the reverse direction, with one “C” insertion. This results in the translation of an aberrant protein of 36 amino acids, which contains only the first 35 amino acids of the normal GATA3 protein (HD: homeodomain, Ala: poly-alanines track, TA: transactivation domain, Zn: zinc finger domain) (**B**) Western blot analysis of PHOX2A, PHOX2B and GATA3 TFs in the 2 *PHOX2A*^−/−^ clones and in the SH-SY5Y cell line, Vinculin was used as a loading control. (**C**) Unsupervised clustering analysis using the top 10% of genes with the highest IQR (inter-quantile range) shows that *PHOX2A^−/−^* and *PHOX2B^−/−^* clones resemble the parental noradrenergic SH-SY5Y cell line whereas *GATA3^−/−^* clones are clustered in the same branch as the SH-EP mesenchymal cell line. (**D**) Volcano plot of a differential analysis comparing the 4 *GATA3^−/−^* samples with 5 noradrenergic neuroblastoma cell lines without *MYCN* amplification (CLB-GA, NB-EBc1, SH-SY5Y, SJNB-1 and SK-N-FI). We used a raw p-value <0.05 and a fold-change >2. Gene Ontology analysis using ToppGene (https://toppgene.cchmc.org/enrichment.jsp) performed on the lists of differentially expressed genes in the *GATA3^−/−^* samples compared to the other neuroblastoma cell lines.

**Figure S2:**
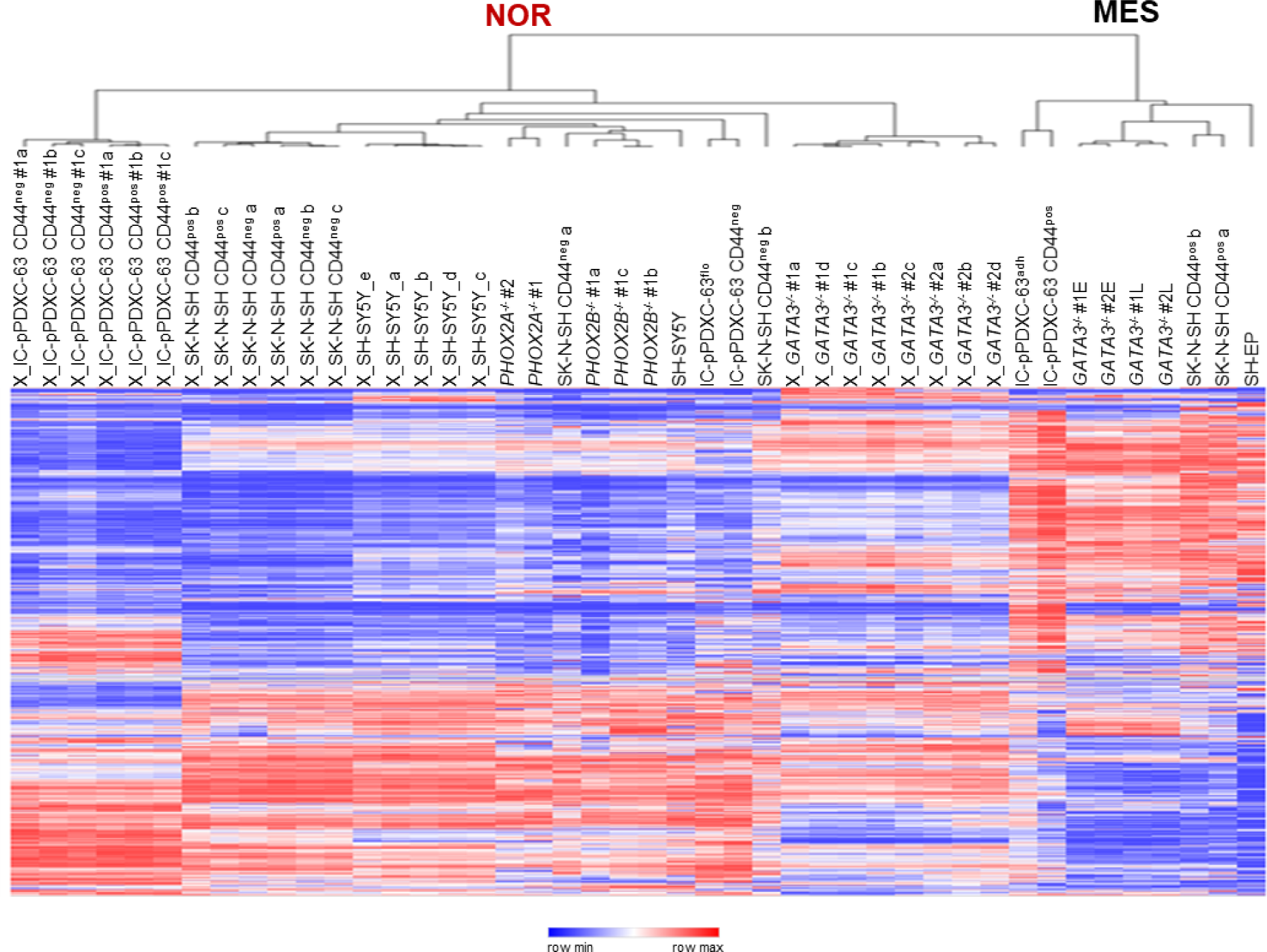
Sorted mesenchymal cells from the IC-pPDXC-63, SK-N-SH and *GATA3^−/−^* cell lines, adopt a noradrenergic identity when engrafted in the mouse. Unsupervised hierarchical clustering based on a transcriptomic signature^13^ discriminating noradrenergic and mesenchymal cells shows that all the *in vitro* mesenchymal cell populations engrafted in mice give tumors with a noradrenergic transcriptomic profile.

**Figure S3:**
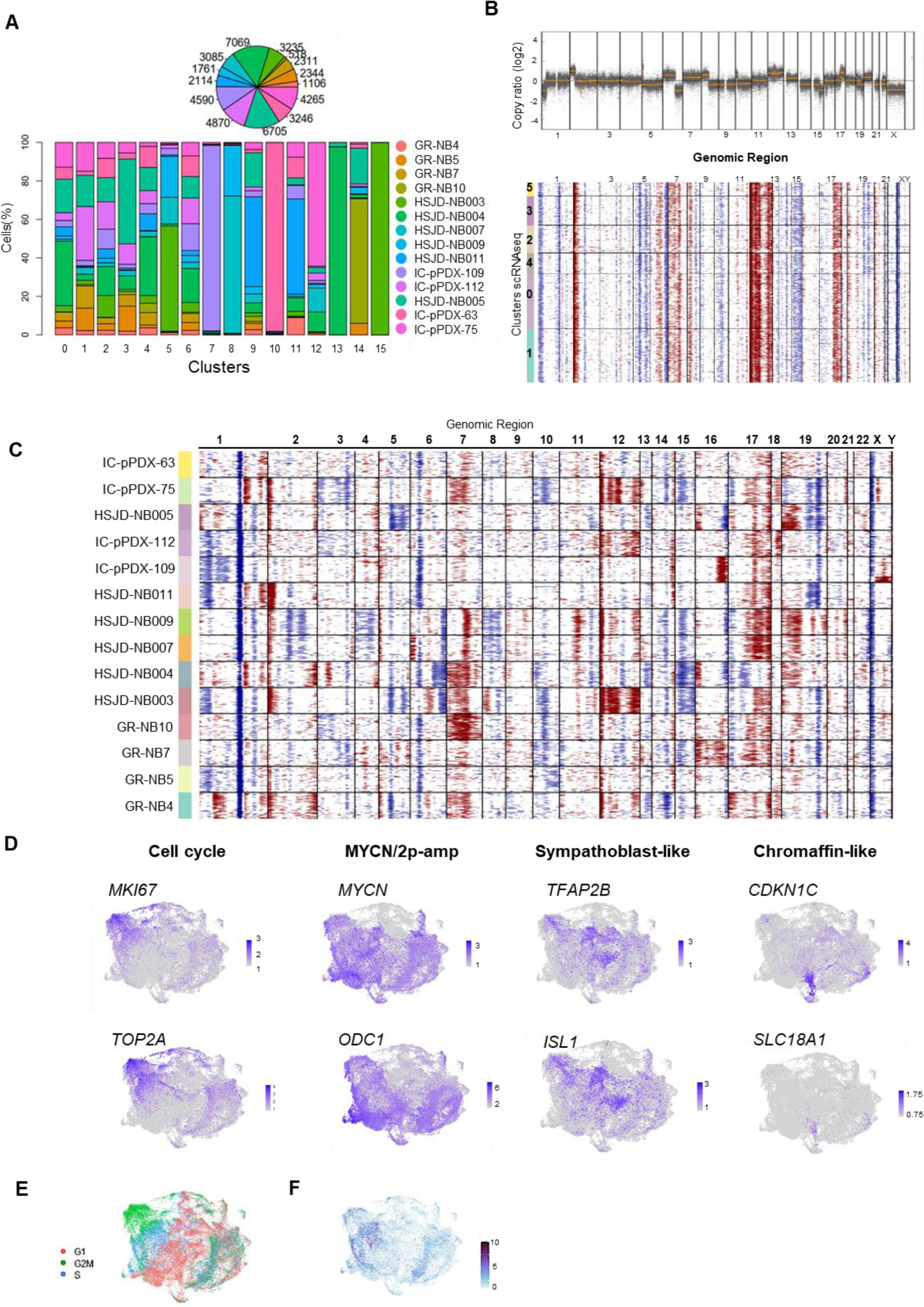
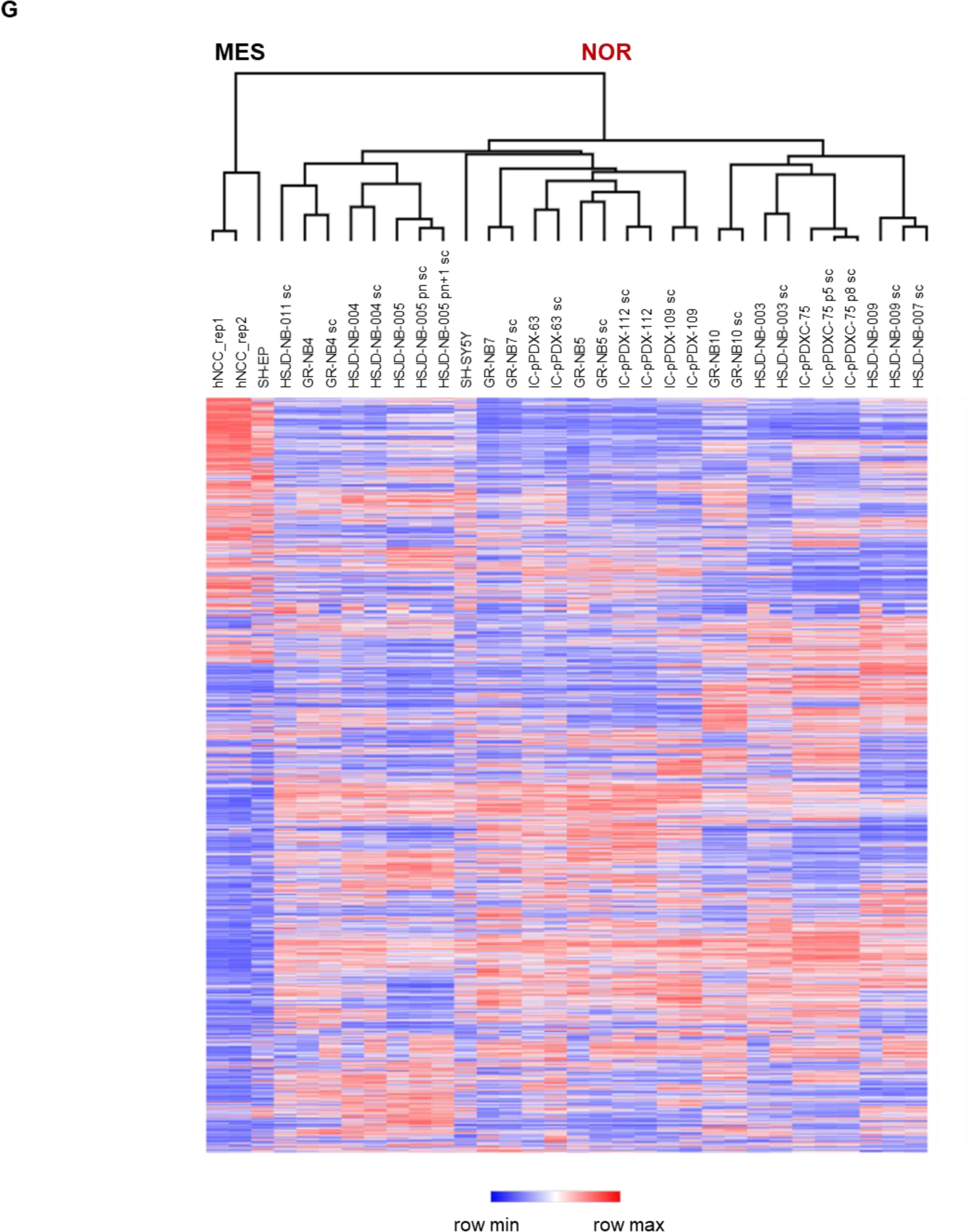
Detailed analyses of single-cell RNAseq of the 14 PDX models. (**A**) The histogram highlights the contribution of the different samples to each cluster in percentage. The pie chart depicts the contribution of each sample to the total number of cells (n= 47,219). (**B**) (Top) Copy number profile inferred from WES from HSJD-NB-003 PDX model. (Bottom) InferCNV profile obtained from scRNAseq data of HSJD-NB-003. (**C**) Overview of the genomic alterations present in each PDX model with the InferCNV profiles on the 14 PDX models, using 500 randomly-selected cells from each sample. (**D**) Individual expression of some genes representing the mains categories identified in the noradrenergic tumor cells: cell cycle, MYCN/2p-amp, sympathoblast-like and chromaffin-like. (**E**) The tumor cells are colored according to their corresponding cell cycle phase (red: G1 phase; green: G2/M phase; blue: S phase). (**F**) Plot of the signature of MYCN target genes^51^ in the integration of all PDX samples. (**G**) Unsupervised hierarchical clustering using the top 10% of genes with the highest IQR (inter-quantile range) for all single cell RNAseq from PDX samples, the mesenchymal (SH-EP and hNCCs) and noradrenergic (SH-SY5Y) cell lines. This shows that all PDX models clustered in the noradrenergic (NOR) branch.

**Table S1:** Expression levels of noradrenergic markers, TFs of the noradrenergic and mesenchymal CRCs in cell lines and xenografts.

**Table S2:** Characteristics of the 14 neuroblastoma PDX models studied by scRNA-seq.

**Table S3:** Filtering of human cells with high quality data for the 14 neuroblastoma PDX models.

**Table S4:** Lists of genes that are up-regulated in the different clusters of noradrenergic cells in a series of 14 neuroblastoma PDX models.

